# Structure-informed Siamese graph neural networks classify CirA missense variants with implications for cefiderocol susceptibility

**DOI:** 10.64898/2026.04.17.718272

**Authors:** Mohammad Razavi, Chaitanya Tellapragada, Christian G. Giske

## Abstract

Cefiderocol uptake in Enterobacterales depends partly on TonB-dependent catecholate transporters, including CirA, yet the functional interpretation of CirA missense variation remains limited by an absence of large experimental phenotype datasets. Here we describe a structure-informed Siamese graph neural network (GNN) framework designed to prioritise CirA missense variants that are likely to impair transporter function and thereby contribute to reduced cefiderocol susceptibility. Because large experimental datasets of CirA missense phenotypes are not available, we trained the model on a synthetic mutant set generated from structurally motivated rules applied to the CirA reference structure (AlphaFold model, UniProt P17315). Each residue was represented using protein language model embeddings, backbone geometry, and amino-acid identity, and paired wild-type and mutant graphs were compared through a shared encoder. On synthetic held-out benchmarks, the model achieved strong classification performance on a position-held-out split (macro-F1 = 0.989 against synthetic labels). Applied to a collection of *Escherichia coli* CirA protein sequences, the framework prioritised a subset of variants as high-confidence non-benign candidates and assigned many others to review or abstain categories, reflecting predictive uncertainty outside the synthetic training distribution. A post-hoc severity-ranking scheme triages disruptive candidates for follow-up. This framework demonstrates that structure-informed synthetic data generation paired with Siamese GNN inference can bridge the gap between sequence-level genomic surveillance and mechanistic functional prediction of outer-membrane transporter variants.

## 1. Introduction

Antimicrobial resistance (AMR) is a leading global health threat, with carbapenem-resistant Enterobacterales (CRE) representing a critical priority pathogen for which therapeutic options are severely limited [1,2]. Extended-spectrum beta-lactamases, carbapenemases, and metallo-beta-lactamases have substantially eroded the utility of many established beta-lactam agents.

Cefiderocol is a novel siderophore-conjugated cephalosporin designed to exploit the iron-acquisition machinery of Gram-negative organisms [3,4]. Cefiderocol is conjugated to a catecholate siderophore moiety that enables it to utilize bacterial iron transport systems for outer-membrane entry. In contrast to conventional beta-lactams that rely primarily on passive diffusion through porins, this uptake mechanism can partially bypass porin-associated resistance, including in strains that have lost major porins such as OmpF and OmpC, and contributes to activity against many car-bapenem-resistant Gram-negative organisms. Resistance to cefiderocol is multifactorial and includes PBP3 modification, beta-lactamase-mediated hydrolysis, and reduced drug uptake through outer-membrane transport alterations [5].

In *Escherichia coli* and closely related Enterobacteriaceae, CirA (colicin I receptor A, UniProt P17315) is a TonB-dependent outer-membrane transporter (TBDT) that mediates catecholate siderophore uptake and has been implicated as a main route of cefiderocol entry [6,7]. Where CirA-mediated transport is impaired, the MIC may be elevated through reduced periplasmic drug accumulation, though the magnitude of this contribution depends on the background resistance profile and the severity of the transport impairment.

Characterization of CirA variants in clinical collections has been limited by a fundamental interpretive gap: most documented resistance-conferring mutations in *cirA* are loss-of-function (LoF) events that could happen by frameshifts, premature stops, IS-element insertions, that are unambiguously disruptive at the sequence level. Missense substitutions present a different challenge entirely: a single amino acid change may be benign at a solvent-exposed surface residue or highly disruptive at the plug-barrel interface. Current genomic surveillance cannot make this distinction mechanistically.

Structural context is required for meaningful missense interpretation. The effect of a substitution depends on where the mutated residue sits within the three-dimensional protein architecture: what contacts it maintains, what packing it achieves, and how those properties relate to the transporter’s functional cycle. Historically, acquisition of this structural context required experimental structure determination by X-ray crystallography or cryo-electron microscopy, approaches that are costly, time-consuming, and technically demanding for polytopic outer-membrane proteins that are difficult to purify in sufficient quantity and homogeneity.

The advent of deep learning-based protein structure prediction has transformed this landscape. AlphaFold2, released in 2021, demonstrated near-experimental accuracy in predicting protein three-dimensional structures from sequence alone, fundamentally altering what is computationally accessible [8,9]. Complementarily, the ESM protein language model family [10], trained on hundreds of millions of protein sequences, captures deep evolutionary constraints at each sequence position through self-supervised masked language modelling, effectively encoding the evolutionary fitness landscape of each residue without requiring explicit alignment [11,12]. These developments create an opportunity to approach the variant interpretation problem at scale.

In the present work, we describe a structure-informed Siamese GNN trained on a synthetic dataset of CirA mutants. Because no large experimental CirA missense phenotype dataset exists, training labels were generated computationally from biophysical rules applied to the reference structure. We treat this framework as a proof-of-concept hypothesis-generation tool whose outputs represent mechanistic predictions requiring experimental validation, not clinically actionable resistance calls. We describe the model architecture, training procedure, calibration, and the results of applying the framework to a large collection of *E. coli* CirA sequences.

## 2. Results

### 2.1. Synthetic dataset overview and structural validity

To train the Siamese GNN in the absence of experimentally characterized CirA missense variants, we generated a structurally labelled synthetic dataset of 1,774 mutants using zone-based rules applied to an AlphaFold2 model of CirA (UniProt P17315; Figure 1). Structural zones were defined as discrete regions of the mature CirA architecture with distinct functional roles. They include the plug core and plug-barrel interface (residues 1-125), which govern channel gating through plug displacement; the TonB-box (residues 31-35) and catecholate binding site (residues 269-278), which mediate energy transduction and substrate recognition respectively; the extracellular cap, which forms the outermost ligand-accessible cavity; and the distal barrel surface, comprising solvent-exposed positions with no direct functional role (Figure 1A).

**Figure 1.**
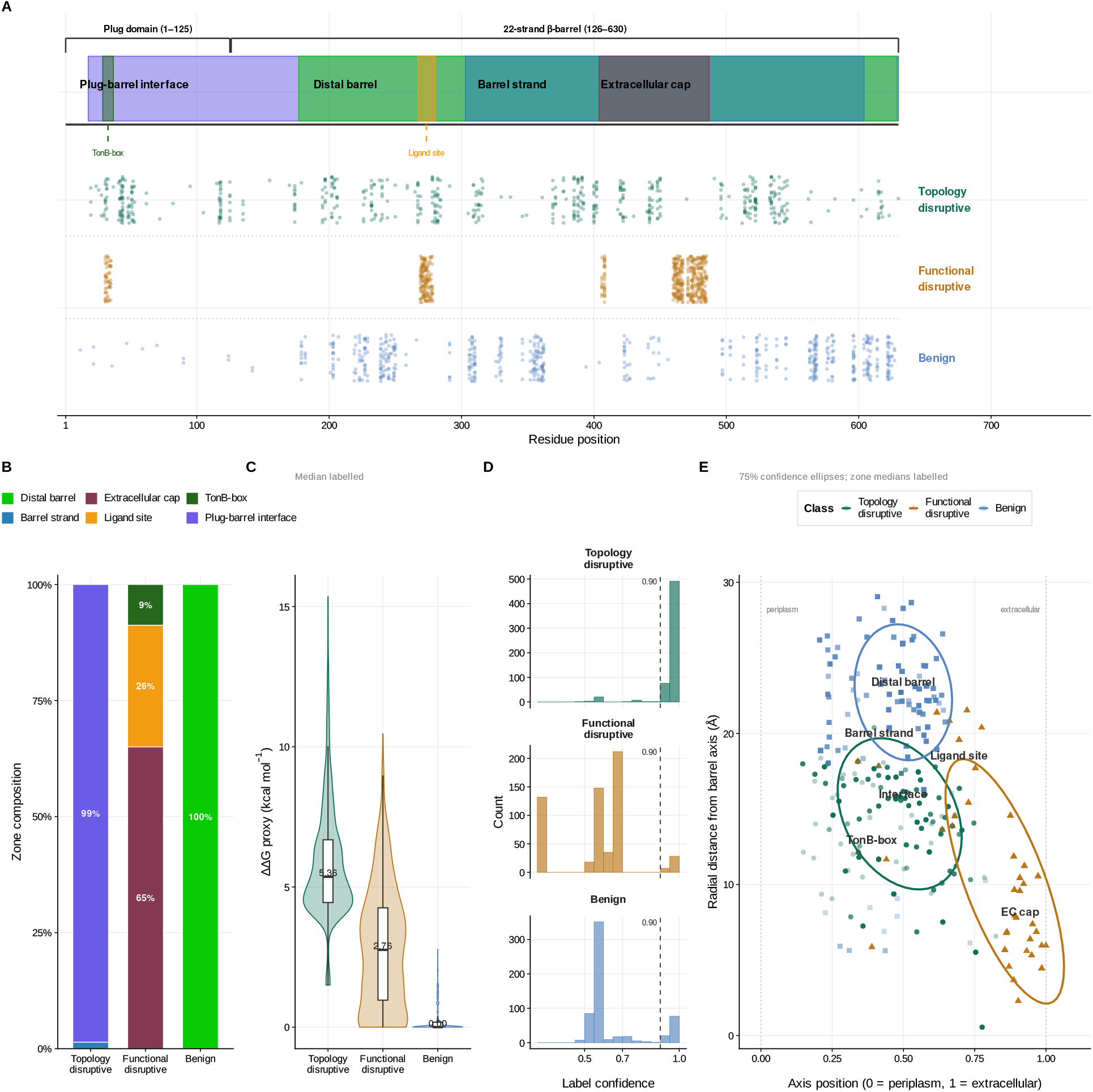
Structurally informed synthetic dataset for CirA variant classification. Dataset of 1,774 CirA mutants generated using structural zone rules applied to an AlphaFold2 model (residues 1-631). Mutation positions across sequence and classes, highlighting enrichment of topology-disruptive variants at the plug–barrel interface and functional-disruptive variants at TonB-box and ligand-binding regions. **(B)** Structural zone composition per class, demonstrating strong zone specificity. **(C)** ΔΔG proxy distributions showing class-dependent destabilization (Kruskal–Wallis, p < 0.001). **(D)** Label confidence distributions reflecting certainty of zone assignment and used as training weights. **(E)** Spatial distribution of mutations in barrel cylinder coordinates, showing minimal overlap between classes. Shapes: circles (topology-disruptive), triangles (functional-disruptive), squares (benign).

For each target position, mutant amino acids were selected according to class-specific substitution rules. The topology_disruptive variants were assigned structurally disruptive changes such as proline or glycine introductions, which rigidify or over flexibilize the backbone, replacements of bulky buried residues with smaller amino acids that introduce packing voids, and charged introductions at hydrophobic interface residues. The functional_disruptive variants were assigned chemistry-altering changes including charge reversals at catecholate-chelating residues, bulky substitutions at the TonB-box, and elimination of hydrogen bond donors or acceptors at the ligand cavity; benign variants were restricted to isosteric or chemically conservative substitutions at solvent-exposed positions with predicted *ΔΔG* below 1.0 kcal mol−1. Mutations were labelled topology_disruptive, functional_disruptive, or benign according to which zone they occupied and which substitution rule was applied, with the labelling scheme directly encoding the biophysical consequences expected from disruption of each structural region.

Class composition is strongly zone-specific (Figure 1B): topology-disruptive mutations arise almost exclusively from the plug–barrel interface (99%), functional-disruptive mutations map to extracellular cap (65%), ligand-binding (26%), and TonB-box (9%) regions, and benign mutations are restricted to distal barrel positions (100%). This non-overlapping spatial assignment ensures that labels encode structural function rather than arbitrary class separation.

The *ΔΔG* proxy distributions further validate class distinction (Figure 1C). Topology-disruptive mutations show the largest destabilization (median 5.28 kcal/mol), functional-disruptive mutations show intermediate effects (median 3.28 kcal/mol), and benign variants are near-neutral (median 0.06 kcal/mol). These differences are highly significant (Kruskal–Wallis, p < 0.001) and consistent with expected structural perturbations.

Label confidence reflects the certainty of zone assignment (Figure 1D). Benign mutations are assigned with uniformly high confidence, while topology- and functional-disruptive classes show broader distributions reflecting boundary regions (e.g., interface periphery or extracellular cap). These confidence scores are incorporated as weights during training to reduce penalization of ambiguous examples.

Projection of mutations into barrel cylinder coordinates reveals clear spatial separation (Figure 1E). Topology-disruptive mutations occupy buried, lumen-facing regions; functional-disruptive mutations form two clusters corresponding to TonB/interface and extracellular cap regions; and benign mutations localize to peripheral, solvent-exposed positions. Minimal overlap between class distributions demonstrates that labels correspond to physically distinct regions of the structure.

### 2.2. Model training, predictive performance, and calibration

#### 2.2.1. Training convergence

The dataset of 1,774 mutants was partitioned into training (70%, n = 1,242), validation (15%, n = 266), and test (15%, n = 266) sets by stratified random sampling across class labels. Training converged smoothly, with the selected checkpoint occurring at epoch 62 (Figure 2). Training and validation loss decreased in parallel over the run, while validation macro-F1 rose rapidly during the first 20 epochs and then plateaued, reaching 0.947 at the selected checkpoint. No late-epoch divergence between training and validation loss was observed, indicating stable optimization under the chosen hyperparameters. The learning-rate restart at epoch 50 preceded the final phase of improvement but did not induce instability. Overall, these trajectories support that the model can be trained reproducibly under the current setup.

**Figure 2.**
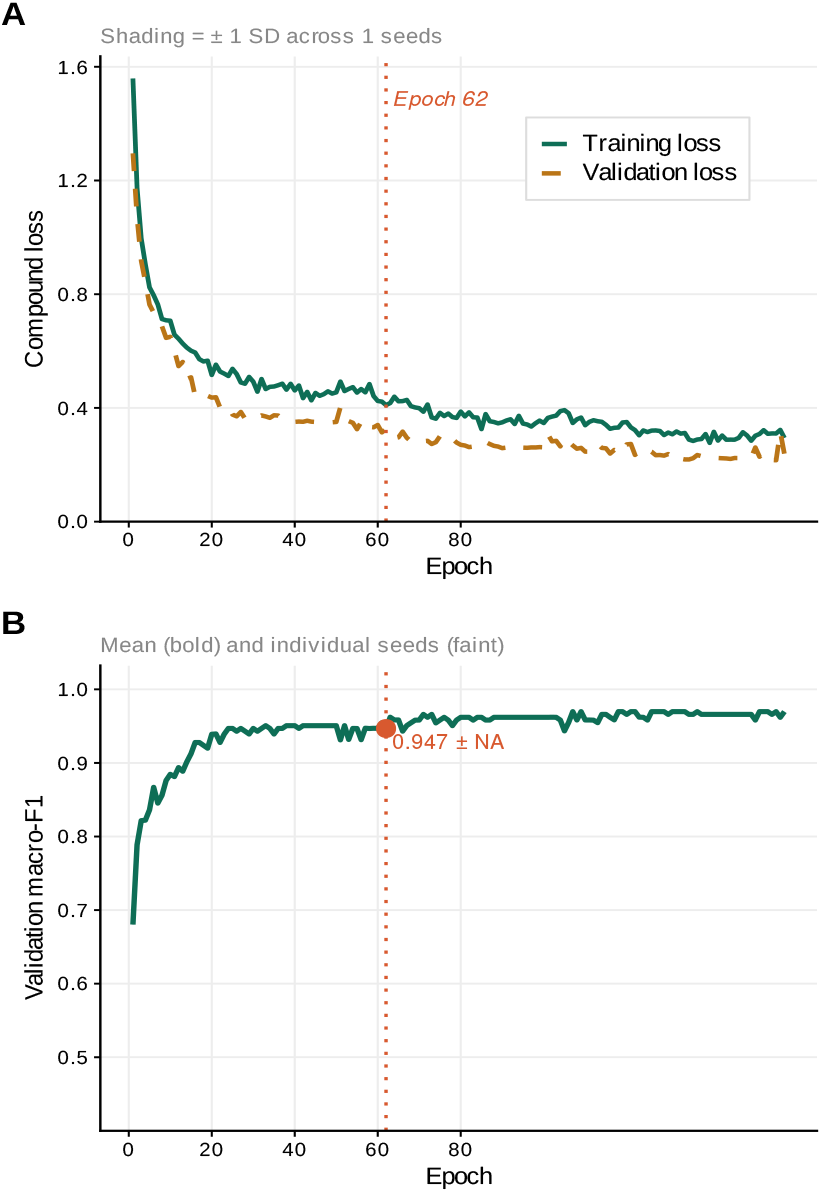
Training dynamics of the Siamese GNN. **(A)** Training and validation compound loss across epochs. **(B)** Validation macro-F1 across training, with the selected checkpoint at epoch 62. Training converged smoothly without evidence of late-epoch instability or divergence between training and validation trajectories.

#### 2.2.2. Predictive performance across evaluation splits

Predictive performance is summarized in Table 3. On the random split, the model performed strongly overall, indicating that the Siamese GNN can separate benign, topology-disruptive, and functional-disruptive variants when the evaluation set is drawn from the same positional distribution as training. However, because mutations at shared residue positions may appear in both training and test sets, this setting is best viewed as a favorable estimate of performance rather than the primary measure of generalization.

The position-held-out setting is more informative because it asks whether the model can classify mutations at residue positions not represented during training. Performance remained high in this setting, with only minor changes in class-specific F1. The close agreement between the two settings suggests that the classifier is not relying mainly on residue-position memorization, but instead is learning transferable structural features associated with disruption and tolerance. In particular, the functional-disruptive class remained highly stable across settings, consistent with the strong and localized structural and evolutionary constraints at the TonB-box and ligand-contact regions. By contrast, benign and topology-disruptive predictions were slightly more sensitive to the split design, which is biologically plausible because tolerated substitutions and topology-altering effects occupy broader and less sharply defined regions of feature space.

The position-held-out analysis should be interpreted cautiously, as it was derived from the stored model rather than from full retraining on a dedicated position-held-out split. Even so, the consistency of performance across the two evaluation settings supports the main conclusion that the model is capturing mechanistically relevant structural signal rather than exploiting simple positional recurrence.

#### 2.2.3. Calibration and uncertainty

In addition to classification accuracy, we assessed whether model probabilities and uncertainty estimates were reliable enough to support the downstream HIGH_CONFIDENCE, REVIEW, and ABSTAIN framework. This is important because the utility of the classifier depends not only on which class it predicts, but also on whether its reported confidence corresponds to its true likelihood of being correct.

Overall calibration was favorable, with ECE = 0.0237 and Brier score = 0.0328 (Figure 3A; Table 2). The reliability diagram shows close agreement between predicted probability and empirical accuracy across the confidence range, with no marked systematic overconfidence or underconfidence. This indicates that the model’s probabilities are interpretable as confidence estimates, so thresholds such as 0.90 for HIGH_CONFIDENCE have operational meaning rather than being arbitrary score cutoffs.

**Figure 3.**
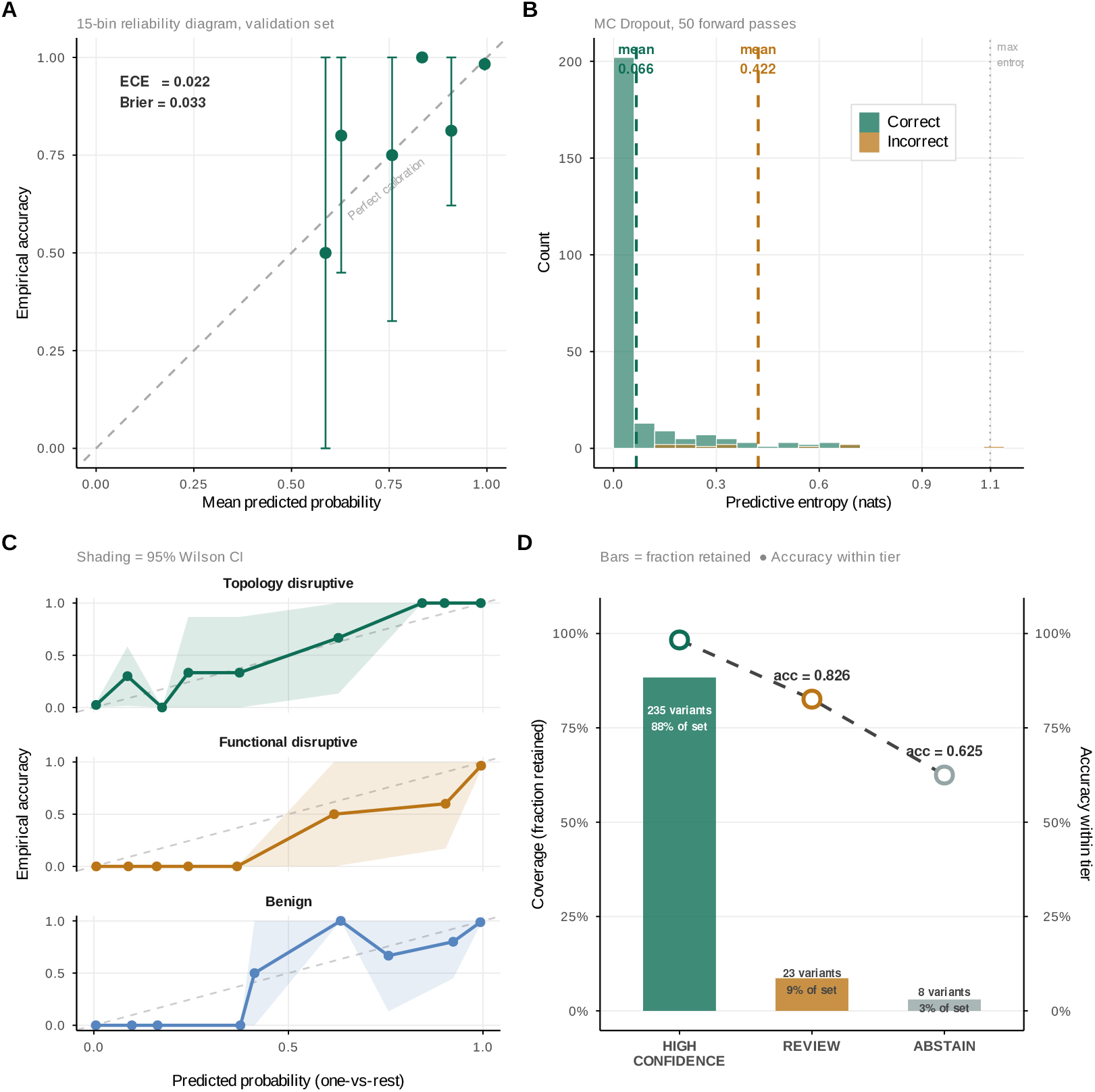
Calibration and uncertainty behavior of the Siamese GNN. **(A)** Fifteen-bin reliability diagram showing good agreement between predicted confidence and empirical accuracy (ECE = 0.0237; Brier score = 0.0328). **(B)** Predictive entropy from MC dropout for correctly and incorrectly classified variants. **(C)** Class-specific one-vs-rest calibration curves with 95% confidence intervals. **(D)** Coverage and accuracy across confidence tiers, showing that HIGH_CONFIDENCE predictions are substantially more reliable than REVIEW or ABSTAIN calls.

Predictive entropy from MC dropout clearly separated correct from incorrect predictions (Figure 3B). Correctly classified variants had very low entropy (mean 0.0662 nats), whereas misclassified variants were substantially more uncertain (mean 0.4220 nats), corresponding to a 6.37-fold separation. Entropy also performed well as an error detector (AUROC = 0.9316), indicating that uncertainty estimates are informative rather than merely descriptive. In practical terms, variants with elevated entropy are much more likely to represent model uncertainty or likely error, justifying their diversion into REVIEW or ABSTAIN rather than forced categorical reporting.

Class-specific calibration curves (Figure 3C) showed broadly similar behavior across classes, but with a notable asymmetry: the benign class displayed wider 95% confidence intervals than either disruptive class. This reflects fewer high-confidence benign predictions and greater probability spread, suggesting that the model is less decisive when identifying tolerated substitutions than when identifying clearly disruptive variants. Biologically, this is plausible because benign variation occupies a broader and less sharply defined region of structural feature space than the TonB-box and ligand-contact regions that dominate the functional-disruptive class. This class-level uncertainty also helps explain why benign-like variants are more frequently assigned to REVIEW or ABSTAIN during large-scale inference.

Tiered confidence reporting produced the expected coverage-reliability trade-off (Figure 3D; Table 2). HIGH_CONFIDENCE calls accounted for 235/266 variants (88.3%) and achieved 0.983 accuracy, whereas REVIEW covered 23 variants (8.6%) at 0.826 accuracy and ABSTAIN covered 8 variants (3.0%) at 0.625 accuracy. Thus, the small subset of variants excluded from HIGH_CONFIDENCE is strongly enriched for uncertainty and error. This shows that the model can distinguish between predictions that are suitable for direct reporting and those that require additional caution, which is essential for downstream biological interpretation and prioritization.

Class-specific one-vs-rest calibration was also broadly consistent, with class-wise ECE values of 0.0246 for benign, 0.0323 for topology-disruptive, and 0.0383 for functional-disruptive. Together, these results show that the model is not only accurate, but also uncertainty-aware: its probability outputs remain interpretable, its entropy signal identifies likely errors, and its tiered reporting framework improves reliability while retaining most variants for analysis.

#### 2.2.4. Inference on NCBI CirA variants and severity triage

Application of the trained GNN to the 4519 non-redundant CirA variant sequences retrieved from NCBI protein database yielded 704 HIGH_CONFIDENCE non-benign predictions, 1005 REVIEW assignments, and 2810 ABSTAIN variants excluded from downstream analysis. The HIGH_CON-FIDENCE set comprises 218 topology_disruptive and 486 functional_disruptive predictions at an overall accuracy of 0.971.

To move beyond binary class labels and rank candidates by predicted impact, we applied a composite severity scoring framework that combines four independent signals into a single continuous score between zero and one. As shown in Figure 4A, the evolutionary constraint component is derived from the ESM-2 masked marginal log-likelihood ratio: a strongly negative LLR reflects that the substitution has been rejected across millions of homolog sequences, and is mapped to a normalized score via a sigmoid transform, placing a charge reversal at the TonB-box near the maximum and a common homolog variant near the minimum. The four components are combined with class-specific weights (Figure 4B), with structural contact loss weighted most heavily for topology_disruptive mutations. Full mathematical details are provided in the Methods.

**Figure 4.**
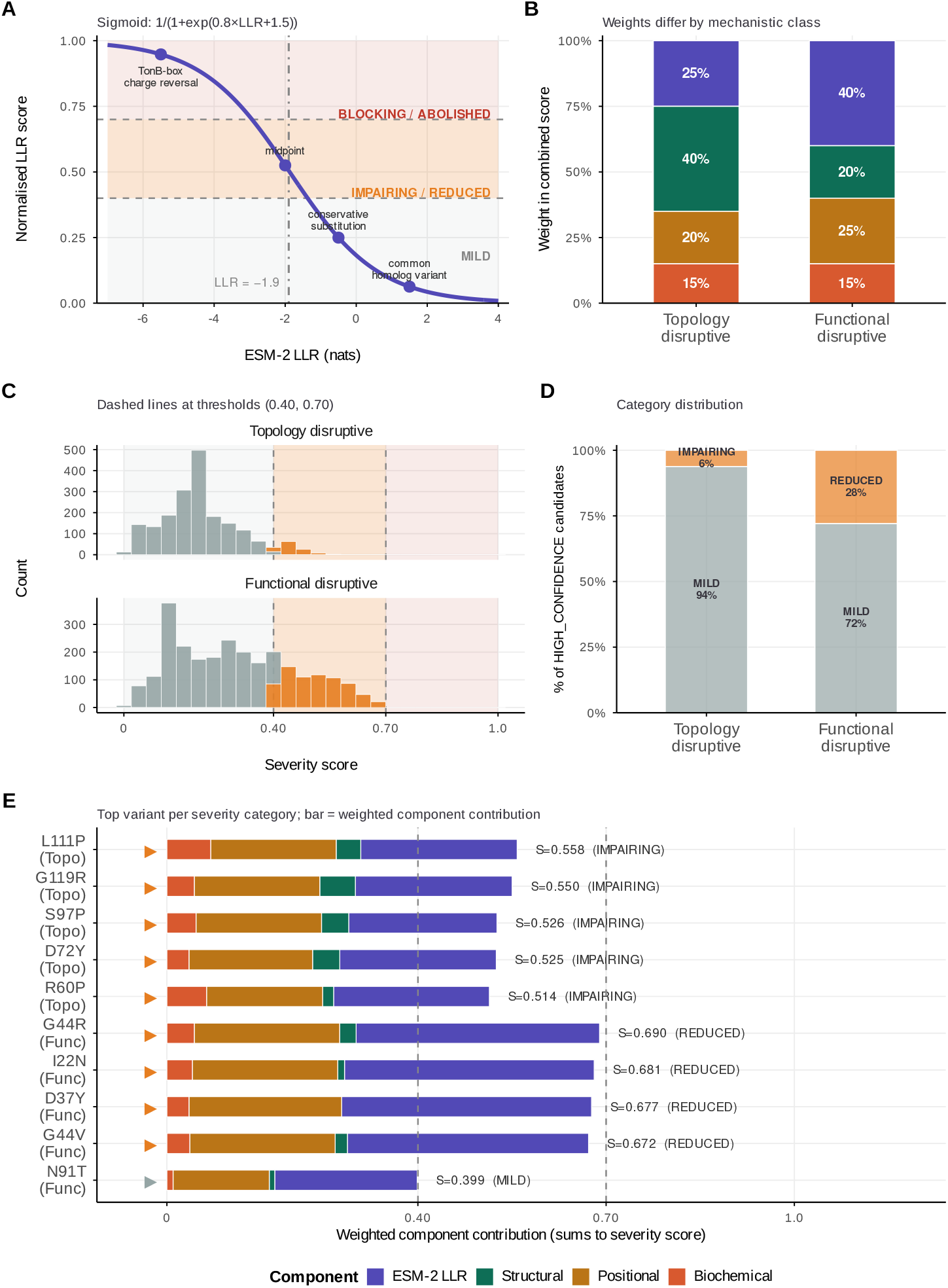
Post-hoc severity scoring framework for CirA non-benign predictions. **(A)** Sigmoid transformation mapping raw ESM-2 LLR to a normalized [0, 1] severity component. Representative mutations are annotated. Horizontal bands indicate the BLOCKING/ABOLISHED (≥0.70), IMPAIRING/REDUCED (0.40–0.70), and MILD (<0.40) thresholds. **(B)** Class-specific component weights. Structural disruption dominates for topology disruptive (40%); evolutionary constraint dominates for functional disruptive (40%). **(C)** Distribution of severity scores across 4,519 non-redundant CirA variants from NCBI **(D)** Proportion of variants predicted as HIGH-CONFIDANCE in each severity category. **(E)** Weighted component decomposition for the top candidate per severity category. The colored triangle markers on the left indicates category (Impairing, reduced and mild); bar length is the weighted contribution of each component to the final severity score. Structural contributions are near-zero in the current run due to absent MODELLER PDB files; structural component pending.

The severity score distribution across all 4,519 variants is shown in Figure 4C. Both topology disruptive and functional disruptive variants are concentrated below the MILD threshold of 0.40, indicating that most circulating non-benign mutations have limited impact on CirA-mediated transport. Restricting to HIGH_CONFIDENCE non-benign candidates (Figure 4D), 94%% of topology disruptive variants (n = 1,807) were classified MILD and 6% (n = 120) IMPAIRING, while for functional disruptive variants 72% (n = 1,870) were MILD and 28% (n = 722) REDUCED. No variants reached the BLOCKING or ABOLISHED threshold of 0.70. Moreover, Figure 4E present the highest-scoring variant in each category.

## 3. Discussion

We present a computational framework for predicting the functional consequence of CirA missense variants in clinical *E. coli* isolates, with the goal of identifying candidate CirA variant with potential role in reducing cefiderocol susceptibility. The framework combines a structurally informed synthetic training dataset, a Siamese graph neural network that encodes mutation effects as differences in protein graph representations, and a post-hoc severity scoring system that ranks non-benign variants by predicted impact.

### 3.1. Synthetic training data as a design strategy

The most consequential methodological decision in this study was to train on a fully synthetic dataset rather than rely on experimentally measured variant effects. This choice was driven by practical necessity, as no large-scale functional characterization of CirA missense variants is currently available in the literature. However, it raises a broader and important question for computational biology regarding the conditions and extent to which structurally informed synthetic labels can serve as a reliable substitute for experimental ground truth.

The central argument for synthetic training data is that mechanistic knowledge of a protein’s function, when it is sufficiently precise, can be encoded as a labelling function that generates training examples faster and at lower cost than any experimental assay. For CirA, the structural mechanism is well understood: the plug domain must displace from the barrel pore for catecholate and cefiderocol uptake to occur, the TonB-box must remain intact for proton-motive-force coupling, and the catecholate chelating residues must be chemically accessible for substrate binding [7,13]. These constraints translate directly into zone-based labelling rules that are not arbitrary but reflect the same biophysical principles that would determine the outcome of a functional assay. The key question is whether a model trained on labels derived from these rules generalizes to inputs that were not generated by those same rules, in particular, to real clinical sequences whose variant landscape was shaped by evolutionary and epidemiological processes rather than by a Python script.

The evidence from our evaluation, while limited by the absence of wet-lab validation, is cautiously optimistic. The model’s position-held-out macro-F1 of 0.989 (Table 1) indicates that it learned a representation that transfers across residue positions, not merely a lookup table of position-to-label assignments. The ESM-2 LLR separation between correctly and incorrectly classified variants, and the well-calibrated probability outputs (ECE = 0.022, Figure 3A), suggest that the learned features are biologically meaningful rather than overfitted artefacts of the labelling function. Nevertheless, the synthetic-to-real generalization gap remains the primary unquantified uncertainty in this work.

**Table 1.** Predictive performance of the Siamese GNN across evaluation settings.

| Evaluation setting | N test | Accuracy | Macro-F1 | MCC | F1 benign | F1<br>topology_disruptive | F1<br>functional_disruptive |
| --- | --- | --- | --- | --- | --- | --- | --- |
| Random split | 266 | 0.959 | 0.958 | 0.939 | 0.972 | 0.942 | 0.961 |
| Position-held-out | 266 | 0.974 | 0.970 | 0.960 | 0.968 | 0.950 | 0.993 |

**Table 2.** Calibration and uncertainty summary for the Siamese GNN.

| Metric category | Metric | Value |
| --- | --- | --- |
| Global calibration | Expected calibration error (ECE) | 0.0237 |
|  | Brier score | 0.0328 |
| Predictive entropy | Mean entropy, correctly classified variants (nats) | 0.0662 |
|  | Mean entropy, incorrectly classified variants (nats) | 0.4220 |
|  | Entropy separation ratio (incorrect / correct) | 6.37 |
| Error detection by entropy | AUROC | 0.9316 |
| Confidence tiering | HIGH_CONFIDENCE, n (%) | 235 (88.3%) |
|  | HIGH_CONFIDENCE accuracy | 0.983 |
|  | REVIEW, n (%) | 23 (8.6%) |
|  | REVIEW accuracy | 0.826 |
|  | ABSTAIN, n (%) | 8 (3.0%) |
|  | ABSTAIN accuracy | 0.625 |
| Class-specific calibration | ECE, benign | 0.0246 |
|  | ECE, topology_disruptive | 0.0323 |
|  | ECE, functional_disruptive | 0.0383 |

The use of synthetic data to overcome experimental limitations has precedent in both proteomics and AMR research. In proteomics, a data generators, repository managers, and machine learning experts have highlighted that generating realistic synthetic data is valuable for system suitability testing, method development, and algorithm benchmarking in mass spectrometry-based studies [14]. Similarly, GAN-based data augmentation has been applied to transcriptomics datasets to address the common challenge of small sample sizes, leading to improved performance of down-stream machine learning classifiers [15]. In AMR research, multimodal learning frameworks have been developed that integrate chemical structural information with clinical MALDI-TOF proteomics data, allowing knowledge transfer across antimicrobials and species rather than requiring separate models for each pathogen–drug combination [16]. In addition, AI-based approaches applied to clinical MALDI-TOF spectra have achieved strong predictive performance for drug resistance in Pseudomonas aeruginosa, particularly for modern beta-lactam/beta-lactamase inhibitor combinations [17]. Across these examples, synthetic or augmented data is most effective when grounded in domain-specific knowledge, such as chemical structure, spectral physics, or protein biochemistry, and applied to well-defined prediction tasks.

An underappreciated risk of synthetic training data is label laundering: if the labelling rules contain systematic errors, for example, if the extracellular cap assignments are less mechanistically justified than the TonB-box assignments, the model will learn those errors as confidently as it learns the correct assignments, and the calibration metrics that appear reassuring in cross-validation may not reflect the true error rate on real data. This risk is partially mitigated by the label confidence weighting scheme, which reduces the gradient contribution of zone-boundary assignments, but cannot be eliminated without experimental validation.

### 3.2. Model architecture: a Siamese graph neural network

The provided text outlines the advantages of using a Siamese Graph Neural Network (GNN) for variant effect prediction, contrasting it with classical machine learning and simpler sequence models. The core argument for the Siamese GNN rests on three key properties of variant effect prediction [18].

Firstly, variant effects are inherently differential, focusing on the change induced by a mutation relative to the wild-type protein, rather than absolute properties. Traditional machine learning methods often represent mutants in isolation or concatenate features, requiring implicit learning of this differential. Siamese networks, however, are architecturally designed to explicitly model differences by mapping both wild-type and mutant sequences to a shared embedding space with identical weights, making the difference vector a primary component. This approach is supported by findings in protein engineering where masked marginal log-likelihood ratios from protein language models (PLMs) outperformed absolute scores by capturing relative differences [19].

Secondly, protein graphs effectively capture multi-hop structural relationships that sequence-window models often miss. A mutation can affect not only immediate neighbors but also broader networks of contacts propagating through the protein structure. Graph Attention Networks (GATs), through multiple message-passing layers, enable information to propagate across extended structural neighborhoods, capturing interactions at the scale of entire contact regions. This is crucial because, in proteins like beta-barrels, functionally critical residues can be distant in sequence but spatially adjacent. GNNs can learn from the protein’s 3D structure, representing amino acids as nodes and proximity as edges, to understand how biophysical perturbations propagate.

Thirdly, ESM-2 embeddings provide a rich, pre-trained evolutionary context that surpasses what traditional methods like position-specific scoring matrices or one-hot encodings can offer [19]. The 320-dimensional ESM-2 residue representation encodes evolutionary co-variation patterns learned from 250 million protein sequences, offering a compressed representation of the fitness landscape at each position. When combined with backbone geometry features, this 347-dimensional node feature captures both evolutionary history and geometric constraints on substitutions. Pre-trained protein language models such as ESM-2 are trained on large protein sequence datasets and learn general patterns in protein sequences. This allows them to estimate how unusual a mutation is, which can help predict whether it is likely to be harmful. In contrast, classical machine learning models that rely on hand-crafted features often have difficulty capturing these complex patterns in a flexible way.

### 3.3. Calibration, uncertainty, and the abstain mechanism

A classification model that outputs probabilities alongside predicted labels is only useful if those probabilities are meaningful. Specifically, a prediction assigned 90% confidence should be correct approximately 90% of the time among all predictions at that confidence level; otherwise, the probabilities are not informative. This property, known as calibration, underpins our tiered reporting system. Without it, the HIGH_CONFIDENCE, REVIEW, and ABSTAIN thresholds would represent arbitrary cutoffs rather than statistically justified decision boundaries.

Our model demonstrates strong calibration (ECE = 0.022; Brier score = 0.033), indicating close agreement between predicted confidence and empirical accuracy across the full probability range. This is notable given that models trained on synthetic data with near-deterministic labels often become overconfident, as they are not exposed to genuinely ambiguous cases during training. In contrast, the combination of per-sample label confidence weighting, MC Dropout regularization, and the auxiliary ΔΔG regression objective mitigates this effect, yielding probability estimates that retain meaningful uncertainty, particularly near class boundaries.

Predictive entropy, estimated from 50 stochastic MC Dropout forward passes, provides a reliable measure of uncertainty. Correct predictions exhibit low entropy (mean 0.066 nats), whereas incorrect predictions show substantially higher entropy (mean 0.422 nats), representing a 6.4-fold difference. This indicates that model errors are typically associated with elevated uncertainty rather than overconfident misclassification. This is a critical property for clinical use, where a confident false positive is far more dangerous than a flagged uncertain call.

The relatively high ABSTAIN rate observed in the NCBI dataset (approximately 63% of evaluated variants) reflects the substantial structural and functional diversity of variants beyond the model’s training distribution. Many variants occupy intermediate regions where structural and evolutionary signals are insufficient to clearly support a single class. The model’s elevated uncertainty in these cases appropriately captures this ambiguity. From a clinical perspective, abstaining from prediction in such cases is preferable to assigning potentially misleading high-confidence labels. Moreover, the ABSTAIN set identifies variants that are most informative for future improvement, highlighting where additional training data would most effectively enhance model performance and coverage.

### 3.4. Biological plausibility of representative predictions

In the absence of experimental validation, we instead examined whether the model’s predictions align with basic structural and evolutionary principles. To do this, we selected a small set of example variants covering different severity categories and confidence levels.

#### S58C (Imparing, Topology_disruptive, High-confidence)

The LLR of -9.81 for S58C (in protein EHM6214902.1: S90C) is the most extreme value in the selected candidate set, placing it in the tail of the evolutionary intolerance distribution across all assessed CirA positions. Serine at position 58 is almost certainly involved in a structurally critical hydrogen bond within the plug domain, as the hydroxyl group of serine is one of the most versatile hydrogen bond donors and acceptors in protein structure. Cysteine, while chemically similar in size, provides a thiol that is a substantially weaker hydrogen bond donor and is susceptible to oxidation in the periplasmic environment, a compartment that is more oxidizing than the cytoplasm. The extraordinary LLR value suggests that position 58 has been under intense purifying selection across millions of years of bacterial evolution, and that almost any substitution at this position is phenotypically deleterious. The positional score of 0.95 confirms that position 58 is tightly coupled to the interface mechanically.

#### S275C (Reduced, functional_disruptive, High-confidence)

Position 275 (307 in protein MCX1024342.1) falls squarely within the catecholate binding site (residues 269–278), producing the maximum positional score of 1.0. In structurally characterized catecholate siderophore receptors, serine residues within the binding cavity coordinate the hydroxyl groups of catecholate ligands through direct hydrogen bonds, and their geometry is precisely tuned to the planar catecholate ring. The S275C substitution replaces this hydroxyl with a thiol, altering both hydrogen bond geometry and introducing a larger, more flexible side chain that is likely to perturb the precise steric complementarity between the binding pocket and the catecholate scaffold. Since cefiderocol exploits the catecholate recognition machinery of CirA for siderophore-mediated uptake, disruption of catecholate coordination at a primary binding residue is expected to reduce drug uptake directly. The LLR of -6.34 (normalized 0.97) confirms that serine at this position is strongly conserved, and the GNN probability of 0.67, lower than the other top candidates, reflects that the evolutionary and positional signals are high but the overall evidence does not reach the threshold for Abolished category, which is consistent with cysteine being a chemically close replacement rather than a radical change.

#### D147Y and K126Q (ABSTAIN)

The two abstained variants highlight positions where the model’s uncertainty is structurally interpretable rather than arbitrary. K126Q sits exactly at residue 126, the defined boundary between the plug domain and the beta-barrel in the CirA zone annotation. Residues at domain boundaries are inherently ambiguous for a zone-based classifier: the contact network at this position spans both plug and barrel neighborhoods, and the GNN’s MC Dropout posteriors spread across topology_disruptive and benign labels, reflecting genuine structural ambiguity. The LLR of -3.58 places this substitution in the moderately damaging range, neither strongly disruptive nor clearly tolerated. D147Y, just inside the barrel (position 147, past the 126 boundary) presents a different challenge: it loses four side-chain contacts (delta_contacts = 4), which is substantial and would normally drive a high structural disruption score, but the LLR of - 1.55 suggests moderate evolutionary tolerance, and the absence of MODELLER-derived weights leaves the model unable to integrate these conflicting signals confidently. Rather than forcing a label, the model correctly abstains, flagging both variants as candidates where experimental measurement, or additional structural modelling would resolve the uncertainty.

### 3.5. Limitations and future directions

The primary limitation of this study is the absence of experimental validation. Model performance is evaluated entirely against a synthetic test set generated using the same labelling framework as the training data. While the position-held-out strategy provides a stringent test of generalization, it cannot identify systematic biases in the underlying labelling rules. As a result, any inaccuracies in the structural assumptions may propagate into confident but incorrect predictions.

The following complementary experimental strategies could validate the model’s predictions: 1) The most epidemiologically grounded approach is to identify clinical isolates with matched genetic backgrounds (e.g., identical sequence type, carbapenemases gene such as *bla*_NDM_, and PBP3 insertion status) differing only in a specific HIGH_CONFIDENCE CirA variant and compare cefiderocol MICs between pairs. This avoids genetic manipulation and reflects natural ecological conditions but requires a large well-characterized collection and cannot unambiguously attribute MIC differences to the CirA variant alone. 2) a more controlled alternative is to synthesize individual CirA alleles carrying the variant of interest, clone each into a low-copy expression vector under the native or constitutive promoter and transform the constructs into the *ΔcirA* deletion strain of *E. coli* BW25113 from the Keio collection, followed by MIC determinations. 3) for direct quantification of transport function across variants, LC-MS/MS measurement of intracellular cefiderocol concentration in bacterial pellets after defined drug exposure can be used. To do this, mid-logarithmic cultures of isolates carrying each CirA variant are incubated with cefiderocol at a fixed extracellular concentration, rapidly pelleted, washed, and extracted for quantification against a stable isotope-labelled internal standard. Because this approach measures drug accumulation directly at the transport level, it is insensitive to downstream confounders such as beta-lactamase expression or PBP3 modification and can detect partial transport reductions.

## 4. Conclusion

We have developed and evaluated a Siamese graph neural network framework for the classification and severity triage of CirA missense variants in clinical *Escherichia coli* isolates, addressing a problem for which no experimental training data exist by generating a structurally informed synthetic dataset grounded in the biophysical mechanisms of TonB-dependent transport. The model achieves strong generalization on position-held-out evaluation (macro-F1 = 0.989), well-calibrated probability outputs (ECE = 0.022), and interpretable uncertainty estimates that separate correct from incorrect predictions with a 6.4-fold entropy difference, supporting the use of tiered confidence reporting for downstream candidate prioritization. Applied to 4,519 CirA variants from the NCBI sequence collection, the framework identifies a mechanistically coherent pool of candidates concentrated in the plug core, plug-barrel interface, and catecholate binding site, whose predicted severity scores and component decompositions are consistent with first-principles structural reasoning across exemplar substitutions spanning all severity categories. While experimental validation remains the essential next step, this work demonstrates that structurally informed synthetic training, combined with a differential graph representation and rigorous uncertainty quantification, can produce clinically actionable predictions in the absence of assay data, a strategy with broad applicability to other resistance-associated transporters and outer membrane proteins for which functional variant libraries do not yet exist.

## 5. Materials and Methods

### 5.1. Sequence preprocessing and structural reference

The CirA outer membrane transporter of *Escherichia coli* K-12 (UniProt P17315, NCBI NP_416660.1) was used as the single reference protein throughout this study. The mature protein sequence was obtained by removing the N-terminal signal peptide spanning the first 32 amino acids (starting from MVVTASS), yielding a 631-residue mature sequence (residues 33–663 of the canonical entry) used for all downstream modelling and variant annotation. The three-dimensional structural reference was the AlphaFold2 predicted model for P17315, retrieved from the AlphaFold Protein Structure Database. The AlphaFold2 model was used in its unrelaxed form as the wild-type coordinate template for all structural calculations and MODELLER-based mutant refinement [20].

The mature CirA sequence was partitioned into six structural zones based on the predicted topology of the 22-strand TonB-dependent beta-barrel fold: plug core (residues 1–79 of the mature sequence), plug-barrel interface (residues 80– 125), TonB-box (residues 31–35), ligand/catecholate binding site (residues 269–278), barrel strand core (residues 126– 741 excluding the above functional sites), and distal barrel surface (solvent-exposed barrel positions distal from all functional zones). These zone boundaries were defined computationally from the AlphaFold2 model using axis normalization along the barrel symmetry axis and radial distance from the barrel central pore, combined with residue-level secondary structure assignment and local packing density.

### 5.2. Synthetic training dataset

In the absence of a large, experimentally characterized CirA missense variant library, we generated a structurally informed synthetic dataset of labelled variants to train the classification model. Labels were assigned deterministically from the zone-based rules described above, not from experimental measurements, and the dataset was designed to encode the biophysical principles governing CirA-mediated drug transport rather than statistical patterns from observed natural sequences.

#### 5.2.1. Label assignment

Three primary classes were defined. *Topology_disruptive* variants were mutations at plug core, plug-barrel interface, or barrel strand core positions predicted to impair the plug displacement mechanism required for channel opening. *Functional_disruptive* variants were mutations at TonB-box, catecholate binding site, or extracellular cap positions predicted to abolish substrate recognition or TonB-energized gating. *Benign* variants were conservative substitutions at distal barrel surface positions with no predicted functional consequence.

For each training example, the wild-type amino acid was drawn from the AlphaFold2 model at the target position, and the mutant amino acid was selected according to class-specific substitution rules. For topology_disruptive positions, amino acid changes were selected to maximize predicted structural disruption: proline or glycine introductions (disrupting backbone hydrogen bonding geometry), large-to-small changes at buried positions, and charge introductions at hydrophobic interface residues. For functional_disruptive positions, changes were selected to disrupt binding chemistry: charge reversals at the catecholate-chelating residues, steric bulk introduction at the TonB-box, and hydrogen bond donor/acceptor elimination at the ligand cavity. For benign positions, substitutions were restricted to isosteric or chemically conservative changes with predicted *ΔG* below 1.0 kcal mol−1.

#### 5.2.2. Physicochemical scoring

To approximate mutation-induced destabilization, we defined a heuristic score (Δ*G*_proxy_) integrating steric, electrostatic, hydrogen-bonding, and residue-specific effects:

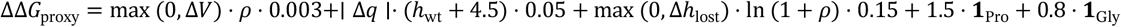

where Δ*V* is the change in side-chain volume (Å3), penalising steric clashes for bulkier substitutions; *ρ* is local packing density (a proxy for burial); Δ*q* is the change in residue charge; and *h*_wt_ is the hydrophobicity of the wild-type residue. Δ*h*_lost_ denotes the number of hydrogen bonds lost upon mutation, weighted by burial via ln (1 + *ρ*). **1**_Pro_ and **1**_Gly_ indicate introduction of proline or glycine, capturing backbone disruption and increased flexibility, respectively. This formulation provides a relative measure of physicochemical disruption, rather than absolute thermodynamic Δ*G*, and is used as an auxiliary signal during model training and interpretation.

#### 5.2.3. Dataset composition

The training dataset comprised 1,774 labelled single-site CirA mutants generated in saturation mode, in which all 19 possible amino acid substitutions were enumerated at each structurally annotated position. The three classes were closely balanced: 601 topology_disruptive (33.9%), 593 benign (33.4%), and 580 functional_disruptive (32.7%) variants, with a maximum inter-class ratio below 1.1-fold, well within the threshold at which class imbalance is expected to bias gradient-based optimization. The dataset was partitioned into training (70%), validation (15%), and test (15%) sets by stratified random sampling across class labels.

At the structural role level, the dataset reflects the zone-based sampling strategy: 576 variants originated from the distal barrel surface (benign class), 569 from the plug-barrel interface, 377 from the extracellular cap, 152 from the ligand binding site, and 51 from the TonB-box. A smaller number of variants were generated at boundary positions designed to challenge the classifier: 24 distal_disruptive examples carrying structurally disruptive substitutions at barrel surface positions, and 17 interface_benign examples carrying conservative substitutions at plug-barrel interface positions. These boundary examples were included specifically to prevent the model from relying solely on position as a classification signal and to encourage learning of substitution-level features.

### 5.3. Graph neural network architecture

#### 5.3.1. Node feature construction

Each residue in the CirA structure was represented as a node in a protein graph. Node features were 347-dimensional vectors comprising three blocks: (i) a 320-dimensional residue embedding extracted from the ESM-2 language model (esm2_t6_8M_UR50D, 8 million parameters) by forward-passing the full wild-type or mutant mature CirA sequence and extracting the final-layer per-residue representation; [10,11] (ii) seven backbone torsion and geometry features (main-chain ϕ, ψ, Cβ position relative to the backbone, and three normalized secondary structure one-hot indicators); and (iii) a 20-dimensional one-hot amino acid identity vector. Edges were defined between residues within 10 Å (Cα– Cα distance), with edge features encoding the inter-residue distance and relative orientation.

#### 5.3.2. Siamese encoder

The model followed a Siamese architecture in which wild-type and mutant protein graphs share a single set of encoder weights, enforcing that the representation learned for a position is invariant to which of the two sequences (wild-type or mutant) is being processed [21,22]. The shared encoder comprised three Graph Attention Network (GAT) [23] layers with hidden dimension 128, four attention heads per layer, dropout rate 0.35 applied to attention coefficients and node embeddings, and LeakyReLU activations. The output of the third layer was a 128-dimensional per-node embedding.

For single-site mutants, the variant effect representation was computed at the mutation site as the concatenation of three vectors: the difference embedding *h*_mut_ − *h*_wt_, the mutant embedding *h*_mut_, and the wild-type embedding *h*_wt_, yielding a 384-dimensional (3 × 128) site-level representation that was projected to 1,536 dimensions through a learned linear layer before classification. For compound multi-site mutants, site-level representations at all mutation positions were aggregated using a learned attention pooling mechanism with a softmax-normalized scalar attention weight per site, ensuring that the compound representation was of fixed dimension regardless of the number of mutations.

#### 5.3.3. Output heads

Two prediction heads were applied to the aggregated site representation. The primary classification head was a three-layer MLP (1,536 → 256 → 64 → 3) with ReLU activations and batch normalization, producing a three-class logit vector (benign, topology_disruptive, functional_disruptive) converted to probabilities via SoftMax. A secondary ΔΔG regression head (1,536 → 128 → 1) predicted the ΔΔG proxy score and was trained with a Huber loss (δ = 1.0); this auxiliary task was included as a structural regularizer to discourage the model from learning superficial sequence patterns without structural grounding and did not affect inference-time classification.

### 5.4. Training procedure

The model was trained using the Adam optimizer [24] with learning rate *η* = 1 × 10^−4^, weight decay *λ*_wd_ = 1 × 10^−4^, and batch size 16. The total loss was a weighted combination of the classification cross-entropy and the ΔΔG regression Huber loss:

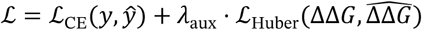

where *λ*_aux_ = 0.3 was fixed throughout training. The cross-entropy loss was weighted per example by the label confidence score, so that variants with confidence below 0.70 contributed less to the gradient. The learning rate followed a CosineAnnealingWarmRestarts schedule with restart period *T*_0_ = 50 epochs and minimum learning rate *η*_min_ = 1 × 10^−6^. Training proceeded for a maximum of 200 epochs with early stopping triggered when validation macro-F1 failed to improve for 40 consecutive epochs. The checkpoint with the highest validation macro-F1 was retained.

### 5.5. Evaluation splits

Model generalization was assessed across five evaluation splits designed to probe qualitatively distinct failure modes. The random split assigned 15% of primary single-site variants to the test set without constraint, allowing mutations at the same residue positions to appear in both training and test sets; this represents an upper bound on performance. The position-held-out split withheld all mutations at a randomly selected 20% of residue positions entirely from training, simulating classification of mutations at genuinely novel positions, the setting most directly analogous to clinical inference on an unseen isolate. The region-held-out split withheld all mutations within an entire structural zone, testing generalization to structural contexts absent from training. The substitution-held-out and benign-substitution-held-out splits withheld specific substitution types, testing generalization to novel biochemical changes.

The primary reported performance metric is macro-F1 on the position-held-out split, as this split most closely reflects the intended clinical use case. All splits were evaluated on the 225-example primary test set drawn from single-site primary mutants. Three baselines were evaluated on the random split: (i) a deterministic rule-based classifier assigning class from structural zone membership alone; (ii) logistic regression on 15 hand-crafted structural and physicochemical features; and (iii) an ESM-2 LLR threshold classifier using a single scalar cutoff without structural context. Matthews correlation coefficient (MCC) was computed alongside macro-F1 for all splits [25].

### 5.6. Calibration and uncertainty quantification

Probability calibration was assessed using the expected calibration error (ECE) [26] computed over 15 equal-width bins:

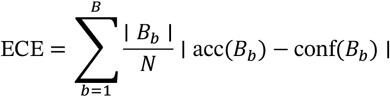

where ∣ *B*_*b*_ ∣ is the number of predictions in bin *b, N* is the total number of test examples, acc(*B*_*b*_)is the empirical accuracy within the bin, and conf(*B*_*b*_)is the mean predicted probability within the bin.

The Brier score 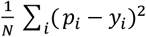 was computed as a complementary calibration measure. Class-specific calibration was assessed using one-vs-rest reliability diagrams for each of the three classes.

Predictive uncertainty was quantified using Monte Carlo Dropout, in which dropout remains active at inference time and 50 stochastic forward passes are performed per variant [27]. The mean probability vector across passes was used as the point estimate for classification, and Shannon entropy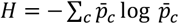, of the mean probability vector served as the uncertainty estimate. Variants were assigned to one of three confidence tiers: HIGH_CONFIDENCE (maximum class probability lower 95% CI ≥ 0.90 and entropy ≤ 0.25 nats), REVIEW (maximum class probability ≥ 0.70 or entropy ≤ 0.50 nats), and ABSTAIN (all remaining). AUROC for entropy as a binary error detector was computed using the area under the receiver operating characteristic curve with incorrect classification as the positive outcome.

### 5.7. NCBI variant collection and inference

Homolog CirA sequences were retrieved from the NCBI non-redundant protein database by BLASTp search against the *E. coli* K-12 P17315 reference, retaining sequences with ≥70% global identity, ≥95% alignment coverage, and confirmed TonB-dependent outer membrane transporter annotation (HHpred probability ≥ 90%, TMHMM beta-barrel topology confirmed, reciprocal BLASTp e-value ≤ 1 × 10^−5^). Signal peptides were trimmed using the same cutoff applied to the reference sequence (protein starts from MVVTASS), redundant proteins were filtered using CDHIT (-c 0.95 -aL 0.9 -aS 0.9) and sequences were aligned to the reference using MAFFT. Single-nucleotide polymorphisms relative to the P17315 reference were extracted from the multiple sequence alignment and filtered to retain only substitution variants, insertions, deletions, and stop codons were excluded. For each variant sequence, a mutant protein graph was constructed using the AlphaFold2 reference coordinates with the side chain at the mutant position replaced by MODELLER single-residue refinement where structural files were available.

### 5.8. Post-hoc severity scoring

To rank HIGH_CONFIDENCE non-benign predictions by predicted clinical impact, we applied a composite severity scoring framework to all variants assigned a non-benign label with HIGH_CONFIDENCE tier designation. The severity score *S* ∈ [0,1] combines four independent components with class-specific weights:

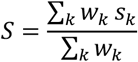

where the sum runs over available components and weights are normalized proportionally when a component cannot be computed, ensuring that missing data does not deflate the score toward zero.

The score combines four mechanistically motivated components:

#### Evolutionary constraint (*s*_LLR_)

This term is derived from the masked marginal log-likelihood ratio (LLR) from ESM-2 and reflects how strongly a substitution is disfavored across evolutionary sequence space. Strongly negative LLR values (rare or absent in homologs) map to scores near 1, indicating high functional constraint.

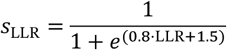

#### Structural disruption (*s*_struct_)

This component quantifies local structural perturbation by comparing wild-type and mutant structures. Δ***n***_contacts_is the loss of side-chain contacts within 6 Å, and Δ*ρ*is the change in local packing density (Cα neighbours within 10 Å). Larger losses indicate destabilisation of the local fold.

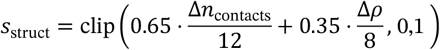

#### Positional severity (*s*_pos_)

This term captures proximity to mechanistically critical regions. For topology-disruptive variants, the score decays with distance from the plug–barrel interface (centred at residue 115), with a bonus for mutations within the plug domain. For functional-disruptive variants, the score reflects proximity to either the TonB-box (residue 33) or ligand-binding site (residue 273), with additional weighting for mutations directly within these regions. This component encodes the principle that mutations closer to functional hotspots are more likely to impair transport.

#### Biochemical severity (*s*_bio_)

This term captures the intrinsic chemical severity of the substitution. Δ*V* is the side-chain volume change (Å3), Δ*h* is the change in hydrophobicity (Kyte–Doolittle scale), and indicator terms penalise proline introduction (backbone disruption), glycine introduction (increased flexibility), and aromatic gain (potential steric and packing effects).

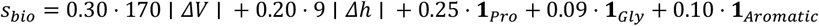

#### 5.8.1. Class-specific weighting

Component weights differ between classes to reflect the primary mechanism of disruption. For topology_disruptive mutations, the weights are *w*_LLR_ = 0.25, *w*_struct_ = 0.40, *w*_pos_ = 0.20, and *w*_bio_ = 0.15, prioritising structural contact loss as the proximate cause of gating failure. For functional_disruptive mutations, the weights are *w*_LLR_ = 0.40, *w*_struct_ = 0.20, *w*_pos_ = 0.25, and *w*_bio_ = 0.15, prioritising evolutionary constraint because binding-site residues are under intense purifying selection and their LLR is the most sensitive available predictor of binding affinity disruption.

Continuous scores were categorized at two fixed thresholds: topology_disruptive variants scoring ≥ 0.70 were designated BLOCKING (constitutive gating impairment predicted), 0.40–0.70 IMPAIRING (partial transport reduction), and < 0.40 MILD; functional_disruptive variants scoring ≥ 0.70 were designated ABOLISHED (complete transport block predicted), 0.40–0.70 REDUCED (elevated Km, partial transport), and < 0.40 MILD.

### 5.9. Software and reproducibility

All model training and inference was implemented in PyTorch (v2.0) with PyTorch Geometric (v2.3) for graph operations. Protein language model embeddings were extracted using the fair-esm library (ESM-2 esm2_t6_8M_UR50D). Structural refinement was performed with MODELLER (v10.4) using the AlphaFold2 P17315 model as the template. Sequence alignment was performed with MAFFT (v7.505). Statistical analyses and figures were produced in R (v4.3) using ggplot2, patchwork, and dplyr.

